# How population control of pests is modulated by density dependence: The perspective of genetic biocontrol

**DOI:** 10.1101/2024.11.08.622719

**Authors:** C. D. Butler, A. L. Lloyd

## Abstract

Managing pest species relies critically on mechanisms that regulate population dynamics, particularly those factors that change with population size. These density-dependent factors can help or hinder control efforts and are especially relevant considering recent advances in genetic techniques that allow for precise manipulation of the timing and sex-specificity of a control. Despite this importance, density dependence is often poorly characterized owing to limited data and an incomplete understanding of developmental ecology. To address this issue, we construct and analyze a mathematical model of a pest population with a general control under a wide range of density dependence scenarios. Using this model, we investigate how control performance is affected by the strength of density dependence. By modifying the timing and sex-specificity of the control, we tailor our analysis to simulate different pest control strategies, including conventional and genetic biocontrol methods. We pay particular attention to the latter as case studies by extending the baseline model to include genetic dynamics. Finally, we clarify past work on the dynamics of mechanistic models with density dependence. As expected, we find substantial differences in control performance for differing strengths of density dependence, with populations exhibiting strong density dependence being most resilient to suppression. However, these results change with the size and timing of the control load, as well as the target sex. Interestingly, we also find that population invasion by certain genetic biocontrol strategies is affected by the strength of density dependence. While the model is parameterized using the life history traits of the yellow fever mosquito, *Aedes aegypti*, the principles developed here apply to many pest species. We conclude by discussing what this means for pest population suppression moving forward.

## 1 Introduction

Insect pest species inflict immense ecological, economic, and medical tolls each year. Perhaps the most notorious pest is the mosquito, a vector for diseases that annually infects hundreds of millions of people. The deadliest of these is malaria and is transmitted by Anopheline mosquitoes. The World Health Organization reports that over 200 million people were infected with malaria in 2022, approximately 600,000 of whom died [75]. Another notable mosquito species, *Aedes aegypti*, vector of many human pathogens, including Zika, Chikungunya, West Nile, and dengue. Half of the world’s population is at risk of exposure to dengue alone [74], and this figure is expected to increase in the coming decades [48]. Control measures for these and other insect pests have typically employed broad-spectrum insecticides. Reducing the pest population alters density-dependent factors, which can hinder or complement control efforts. For example, *Ae. aegypti* populations are regulated by density-dependent juvenile mortality as a result of intraspecific competition [33]. Adding mortality before this competition can mitigate density-dependent regulation and counterintuitively increase the adult population [1, 4, 47]. Effective population control then becomes a matter of best navigating (or exploiting) these natural processes. Understanding how density dependence affects population suppression is vital to this end.

We pay particular attention to *Ae. aegypti* given its medical importance as a vector responsible for significant human morbidity and mortality. As stated above, density dependence in *Ae. aegypti* occurs during the larval stages as a result of resource competition [9, 64, 80]. Insufficient nutrition caused by competition can lengthen the time to or prevent pupation [2, 49, 70, 71, 72, 80], and decrease adult body size [2, 13, 49, 70, 71, 72, 80]. Similar density-dependent factors occur in many organisms and the principles developed here can easily be extended to other pest species. Indeed, DD juvenile regulation is a common trait in holometabolous insects, including many Lepidopteran pests [66]. Beyond this, recent innovations in genetic control methods of *Ae. aegypti* (e.g., [6, 39]) have renewed interest in understanding the impact of density dependence.

One such innovation is gene drive. Gene drives any natural or artificial mechanism of propagating a gene into a target population [3]. The introduced gene propagates in the target population even if the cargo is deleterious to carrier organisms. Gene drives have gained considerable attention recently owing to the success of *in vivo* studies in different pest species, including *Ae. aegypti* [5, 39], the malaria mosquito *Anopheles gambiae* [37], and the crop pest *Drosophila suzukii* [76]. These advances have also made it possible to adjust the sex specificity of a control (e.g., female-specific flightlessness in mosquitoes [22]), as well as the ability to cause lethality at a particular time in the organism’s development.

Much of the research investigating the role of density dependence in population control has focused on a particular strategy or method. For example, for conventional controls, May et al. (1981) found that a parasitoid can increase host abundance depending on the sequence of parasitism and density-dependent (DD) regulation [45]. Some of these early results, however, disagree in their conclusions. In his seminal work using field data of *Ae. aegypti* populations, Dye (1984) constructed several analytical models to simulate mosquito population dynamics [20]. This included a continuous-time model with an exponential form of DD juvenile mortality. His sensitivity analysis revealed that targeting adults was more effective at population control than reducing birth rates or larval breeding sites. In marked contrast to Dye’s findings, Newton and Reiter (1992) considered a simple continuous-time model with logistic growth and a linear growth rate to assess the efficacy of ultra-low volume (ULV) spraying [52]. They found that culling adult mosquito populations via ULV spraying was inferior to reducing the environmental capacity of mosquitoes, such as via source reduction.

Early field trials of the sterile insect technique (SIT) found evidence that DD regulation hindered control success. McDonald et al. (1977) observed that pupal production did not significantly decrease following the release of sterile males [46]. The investigators concluded that this was caused by DD larval mortality and hypothesized that DD changes in larval mortality would work against autocidal control strategies in general. More recent work on SIT and other genetic control measures has reinvigorated the discussion of how density dependence interacts with population suppression. The investigators in [77] found that partial population sterilization can increase the survival of density-regulated larvae. Vella et al. (2021) studied the strength of density dependence alongside the timing of mortality due to control but for repeated releases of mosquitoes carrying a female-killing gene, finding that stronger density dependence required larger release ratios to achieve the same level of suppression [68]. The authors in [19] hypothesized how different strengths of density dependence affect a population’s response to suppression by a gene drive, which we explore in the present manuscript. Both field and modeling studies have demonstrated how specific control outcomes are modulated by density dependence. With these observations in mind, a general treatment of the interaction between density dependence and population control is appropriate.

We construct a baseline continuous-time population model with sex, age, and DD juvenile mortality, similar to the model in [68]. Population control is included in the model using a general control load, the timing of which can be either early-acting or late-acting, and can target both sexes or females exclusively. We ignore male-sex control loads because males tend to be innocuous or incapable of transmitting disease [55]. In addition, for *Ae. aegypti*, males are not reproductively limiting and mate multiple times in their lifetime [30, 32]. Early-acting control loads take place before DD mortality and simulate control strategies that reduce female fecundity. Examples include larvicides, SIT [10], and *Wolbachia* [69]; for the latter two, female fecundity is reduced as a result of mating with sterile or incompatible males. Late-acting control loads are imposed after DD mortality. RIDL (Release of Insects carrying a Dominant Lethal) is an example of a control program for *Ae. aegypti* that utilizes late-acting mortality [67]. Traditional control strategies such as insecticide-treated bed nets and adulticides are also late-acting as only adults are killed. For the baseline model, the equilibrium population size of each system following control—hereby referred to as the control equilibrium—is calculated, as well as the reproduction number. In performing this analysis, we clarify earlier work on density dependence in discrete- and continuous-time population models. The data available on density dependence in *Ae. aegypti* does not lend itself to a single correct mathematical interpretation [38], so any attempt to include it in a model must be accompanied by some amount of guesswork. To account for this, we study different functional forms of juvenile DD mortality distinguished by their strength of density dependence. Our results are made more generalizable in doing so. Finally, the baseline model is extended to consider three gene drives as case studies: homing, one-locus engineered underdominance, and two-locus engineered underdominance. The analysis performed for the baseline model is repeated for each case study. We conclude with the consequences of our findings, including implications for future models.

## 2 Models

### 2.1 Baseline model

We first outline a general ODE model to study the interaction between population suppression and DD juvenile mortality. Let *J*_*i*_ and *A*_*i*_ denote juvenile and adult populations, respectively, with the subscript indicating sex—*M* for males, and *F* for females. The general function *f* (*J*) represents the per capita DD juvenile mortality rate, for *J* = *J*_*M*_ + *J*_*F*_. Our model is given below.

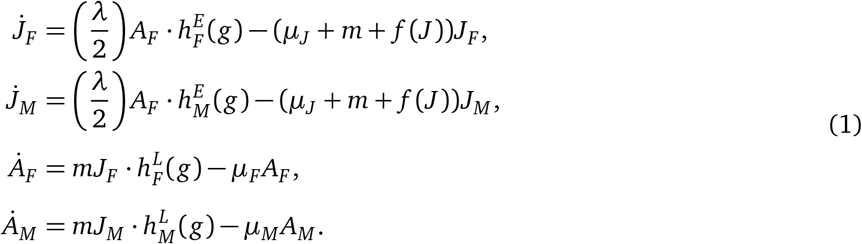

Here, dots denote differentiation with respect to time. Juveniles mature at rate *m*, while density-independent mortality occurs at per capita rate *µ*_*J*_. Female adults produce eggs at a daily rate *λ* and have an average lifespan of 1*/µ*_*F*_, while male adults have an average lifespan of 1*/µ*_*M*_. We assume that reproduction is not male limited and that the number of new offspring depends on *A*_*F*_ only. Definitions and values for parameters, variables, and functions are given in Table 1. Pest mortality due to the control can occur before (early-acting) or after (late-acting) DD juvenile mortality. We refer to the imposed load as the control load, denoted by the variable *g* ∈ [0, 1]. This is the proportion of insects killed at a certain life stage. Inclusion of the control load is done via the functions 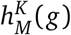 and 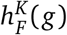 for males and females, respectively. The timing of population suppression is indicated by the superscript *K* ∈ *{E, L}* depending on whether the control load is early-(*E*) or late-acting (*L*). For example, in the case of an early-acting control load that affects both sexes, 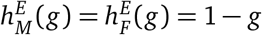 and 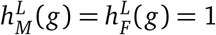.

**Table 1:** Parameter, variable, and function information for the systems in Eqtns. (1) and (2).

| Parameter | Description | Value | Source |
| --- | --- | --- | --- |
| $\lambda$ | Daily female egg-laying rate (per capita) | $8 \text{ d}^{-1}$ | [27, 65] |
| $p_{ij}(t)$ | Probability of genotype $ij$ offspring | varies | N/A |
| $\varphi_{M,ij}, \varphi_{F,ij}$ | Relative fitness of males, females with genotype $ij$ | varies | N/A |
| $h_M^E(g), h_F^E(g)$ | Early-acting control load for males, females | varies | N/A |
| $h_M^L(g), h_F^L(g)$ | Late-acting control load for males, females | varies | N/A |
| $\mu_J$ | Density-independent mortality rate of juveniles (per capita) | $0.029 \text{ d}^{-1}$ | [78] |
| $\mu_F$ | Mortality rate of female adults (per capita) | $0.1 \text{ d}^{-1}$ | [21, 50] |
| $\mu_M$ | Mortality rate of male adults (per capita) | $0.28 \text{ d}^{-1}$ | [21, 50] |
| $m$ | Maturation rate (per capita) | $0.14 \text{ d}^{-1}$ | [50] |
| $c_a, c_t$ | Ambient fitness cost, toxin lethality | varies | N/A |
| $g$ | Control load | varies | N/A |

We consider several functions for the per capita juvenile DD mortality rate, *f* (*J*). The functions we study are motivated by their usage elsewhere in the mosquito modeling literature.

1. Generalized logistic case: *f* (*x*) = *αx*^*β*^, used in [11, 54, 58, 59, 68], among many others. We analyze system performance for *β* = 0.5, *β* = 1, and *β* = 1.5.
2. Logarithmic case: *f* (*x*) = ln(1 + (*αx*) ^*β*^), used in [35, 34], with both of these works citing its usage in [7].

Each function above differs in its strength of density dependence, by which we mean how quickly a perturbed population returns to equilibrium [62]. This definition is useful for comparing the efficacy of population control in systems with different functional forms of density dependence. The return time of the system in Eqtn. (1) for each density dependence function is plotted in Fig. 1. Perturbations are simulated by increasing the adult population by a certain proportion, while the return time is how long it takes for the total population to return to within 10^−4^ relative to its (pre-perturbed) equilibrium value.

**Figure 1:**
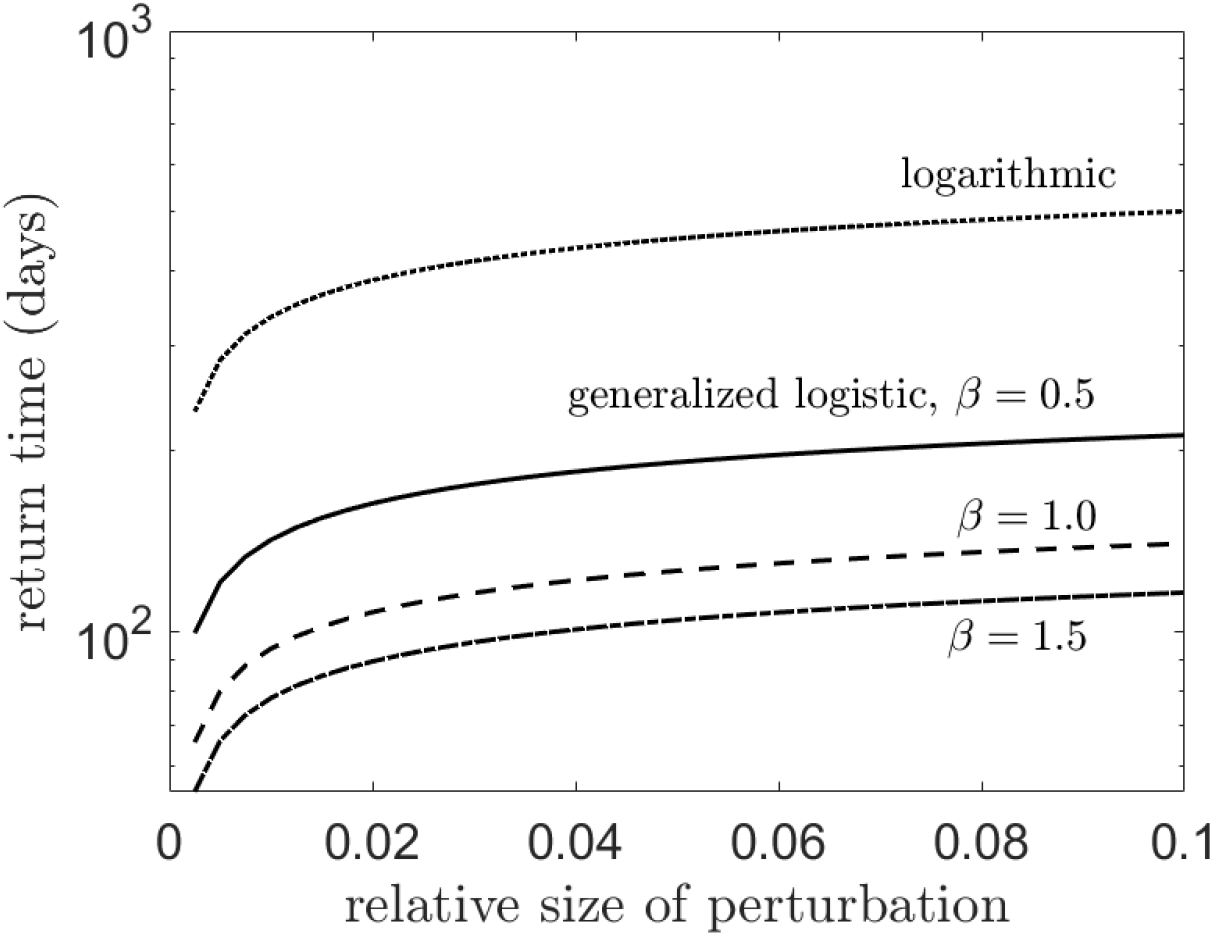
One way to characterize the strength of density dependence is by measuring the time it takes for a perturbed population to return to equilibrium; stronger density dependence produces a shorter return time, while weaker density dependence produces a longer return time. The plot above shows this for different forms of *f* (*J*), or density-dependent juvenile mortality. The generalized logistic case for *β* = 0.5 is given by the solid line, *β* = 1 by the dashed line, *β* = 1.5 by the dash-dotted line, and the logarithmic case by the dotted line.

A few important clarifying points are made in the Appendix about the logarithmic case. Briefly, Bellows [7] derives this equation by integrating a continuous-time function over a single time unit (albeit using a problematic assumption) to arrive at a discrete-time analog that exhibits overcompensatory density dependence.

Consequently, some studies (e.g., [35, 34]) employ the logarithmic form and incorrectly claim that their model exhibits overcompensatory density dependence or scramble competition. A continuous-time formulation of this type is unable to exhibit overcompensatory density regulation, although this does not apply to models that employ a time delay (e.g., [4]). The models used in the present manuscript are continuous-time without time delays so they do not exhibit overcompensatory density dependence.

### 2.2 Genetic model

The mathematical model in Eqtn. (1) is extended with genetic structure to study how genetic control performance depends on the strength of density dependence.

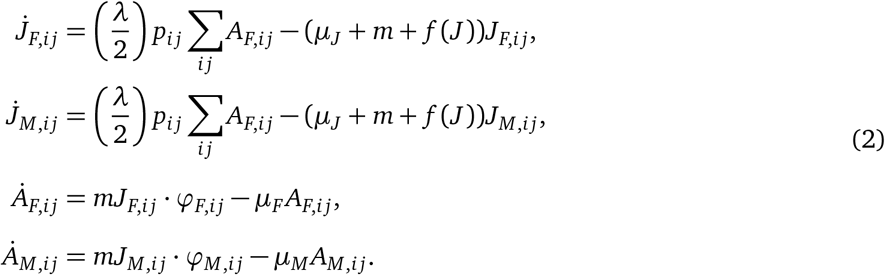

The state variables are the same as those in Eqtn. (1) except now there are subscripts to indicate genotype. The total juvenile population is *J* = Σ_*i j*_ *J*_*M,i j*_ + *J*_*F,i j*_. The probability of an offspring having genotype *i j* at time *t* is given by the function *p*_*ij*_(*t*). For notational brevity, we suppress the dependence on *t*. We let *ϕ*_*M, i j*_ (*ϕ*_*F,i j*_) denote the relative fitness of males (females) with genotype *i j*. We assume these relative fitnesses affect viability after DD competition occurs and are thus synonymous with the late-acting control loads in Eqtn. (1). As in the baseline model, both bi- and female-sex targeting scenarios are considered. All remaining parameters are as they appeared previously. We assume that fitness costs combine multiplicatively.

## 3 Gene drives

As stated above, gene drives are any natural or artificial mechanism of propagating a gene into a target population [3]. Gene drives spread by biasing their inheritance. Gene drives are introduced into wild populations by the release of transgenic, or drive-bearing, organisms. In all of our simulations, we assume that transgenic release occurs once. Transgenic releases can be male-only, female-only, or bi-sex. While some gene drives can spread from arbitrarily low frequencies—such as a simple homing gene drive [15]—for other drives, invasion is a frequency-dependent process. These so-called “threshold drives” require the release of sufficiently many transgenic organisms before the drive can propagate. This figure is quantified as a ratio of released to wild-type organisms. We refer to the ratio required for successful invasion as the invasion threshold. In this work, we study a simple homing drive and two versions of engineered underdominance.

A homing drive biases its inheritance by exploiting DNA repair mechanisms following target allele disruption [15]. In heterozygotes, the transgene targets and cleaves an allele on the homologous chromosome. The break is (ideally) repaired via homology-directed processes, duplicating the transgene and converting the heterozygote into a homozygote. We denote wild-type alleles with Roman letters and transgenic alleles with Greek letters. For a homing drive, there are three genotypes: *AA* (wild-type), *Aα*, and *αα*. We let *c*_*a*_ denote the fitness cost imposed by the transgene. Heterozygote fitness is additionally affected by the dominance of the transgene, denoted by *h. h* = 0 corresponds to a fully recessive transgenic allele, while the fully dominant case occurs for *h* = 1. We take the transgene to be fully recessive, i.e. *h* = 0. Table 2 shows the relative fitness of each genotype for a simple homing drive for both female- and bi-sex targeting scenarios.

**Table 2:** Relative fitness of each genotype for a simple homing gene drive. Here, *s*_*a*_ = 1 − *c*_*a*_.

|  | Genotype |  |  |  |  |  |
| --- | --- | --- | --- | --- | --- | --- |
|  | Male |  |  | Female |  |  |
| Lethality | AA | Aα | $\alpha\alpha$ | AA | Aα | $\alpha\alpha$ |
| Bi-sex targeting | 1 | $s_a(1-h)$ | $s_a^2$ | 1 | $s_a(1-h)$ | $s_a^2$ |
| Female sex targeting | 1 | 1 | 1 | 1 | $s_a$ | $s_a^2$ |

We also study two forms of engineered underdominance (EU) gene drives: one- and two-locus EU. Underdominance refers to when individuals heterozygous for an allele have lower fitness [28]. In the one-locus case, the constructs are carried at the same locus on a homologous pair of chromosomes [18]. Each transgene possesses two main components: an antidote and a toxin. The antidote on one engineered allele rescues the organism from the toxin on the complementary allele. The organism is nonviable if it possesses one transgene without the complement. Let *α* and *β* denote the complementary transgenes (i.e., *α* has the antidote for the *β* toxin and vice versa) so that the genotype of the released transgenic organisms is *αβ*. The genetic payload is twofold: the ambient fitness cost of possessing a transgene, denoted *c*_*a*_, and the lethality of the toxin absent the complementary antidote, denoted *c*_*t*_. The respective relative fitnesses are *s*_*a*_ = 1 − *c*_*a*_ and *s*_*t*_ = 1 − *c*_*t*_. We assume the toxin is lethal to the carrier organism, i.e. *c*_*t*_ = 1, so that the only viable genotypes are *αβ* and wild-type (*AA*). Table 3 shows the relative genotype fitness for female- and bi-sex targeting scenarios in the one-locus EU system.

**Table 3:** Relative fitness of each genotype for a one-locus EU gene drive. Here, *s*_*a*_ = 1 − *c*_*a*_ and *s*_*t*_ = 1 − *c*_*t*_. We take *c*_*t*_ = 1.

|  | Genotype |  |  |  |  |  |  |  |
| --- | --- | --- | --- | --- | --- | --- | --- | --- |
|  | Male |  |  |  | Female |  |  |  |
| Lethality | AA | Aα, Aβ | $\alpha\alpha$, $\beta\beta$ | $\alpha\beta$ | AA | Aα, Aβ | $\alpha\alpha$, $\beta\beta$ | $\alpha\beta$ |
| Bi-sex targeting | 1 | $s_a s_t$ | $s_a^2 s_t^2$ | $s_a^2$ | 1 | $s_a s_t$ | $s_a^2 s_t^2$ | $s_a^2$ |
| Female sex targeting | 1 | $s_a$ | $s_a^2$ | $s_a^2$ | 1 | $s_a s_t$ | $s_a^2 s_t^2$ | $s_a^2$ |

The second form of EU gene drive we consider is the two-locus case. The difference between a two-locus EU and a one-locus EU drive is that for the former, the construct is carried at different loci on non-homologous chromosomes. The main components of each transgene, however, remain the same. We denote the complementary transgenic alleles by *α* and *β* and let *A* and *B* denote the wild-type alleles at either locus (wild-type organisms have genotype *AABB*). There are nine genotypes in total. We assume that an individual needs only a single copy of the complementary transgene to rescue the organism from the toxin. For example, *αAββ* individuals are still viable despite having two copies of the *β* allele and only one copy of the *α* allele. This is sometimes referred to as strongly suppressed underdominance [18]. The viable genotypes are *αAββ, αAβ B, ααββ, ααBβ*, and wild-type (*AABB*). The relative fitness of each genotype for the two-locus EU system is given in Table 4.

**Table 4:**
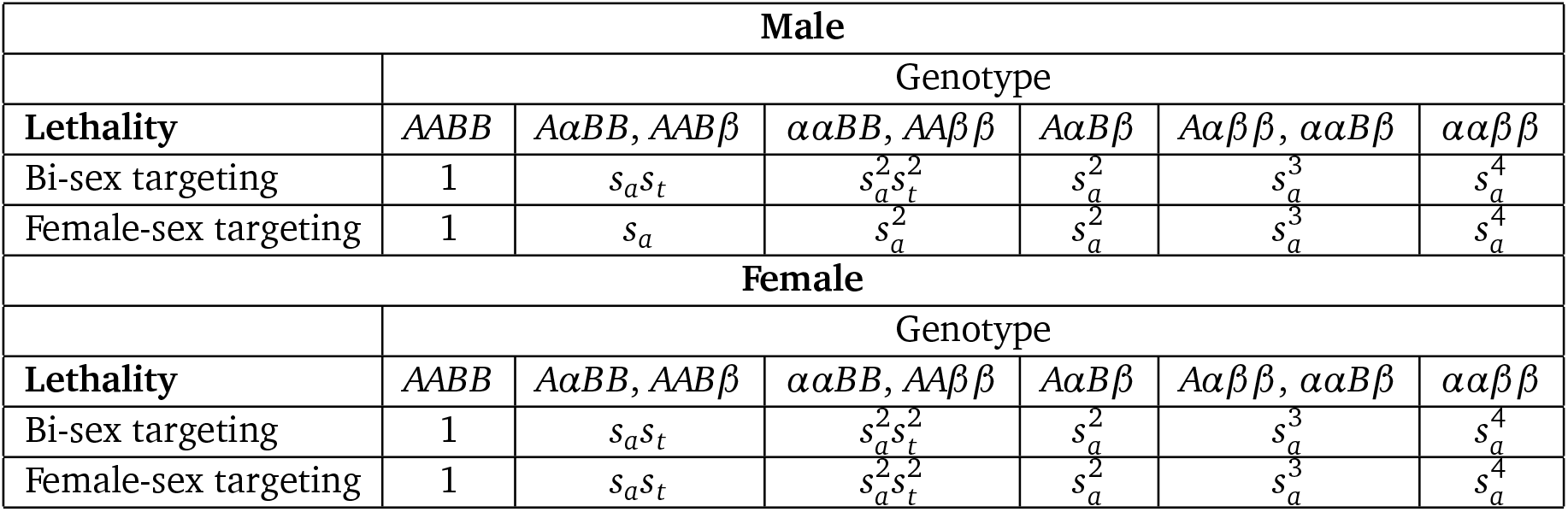
Relative fitness of each genotype for a two-locus EU gene drive with strongly suppressed transgenes. Here, *s*_*a*_ = 1 − *c*_*a*_ and *s*_*t*_ = 1 − *c*_*t*_. We take *c*_*t*_ = 1.

For EU gene drives, the sex ratio of the released population is important, as this affects how efficiently the payload propagates. For a one-locus EU drive, a bi-sex release is required for introgression of the transgene if the toxin is lethal (*c*_*t*_ = 1). This is because all progeny arising from the matings with the released insects are nonviable. Even if the toxin is female-sex targeting, no female progeny will ever be viable and the construct will be unable to sustain itself in the population. Female mosquitoes can serve as disease vectors, making their release undesirable from a regulatory and public health perspective. However, certain genetic control methods additionally render female mosquitoes immune or incapable of transmitting disease [14, 69]. In the two-locus case, we assume that releases are male-only. We study the performance of each of the gene drives discussed above using the model in Eqtn. (2).

## 4 Results: General control load

Control load timing is important and depends on the most harmful life stage of the pest, when population suppression is most effective, and the control mechanism itself. In the following sections, we model an idealized population control strategy in which a control load (*g*) is either early- or late-acting, and either female-or bi-sex targeting. The control load is modeled by assuming a certain proportion of the population is killed.

### 4.1 Early-acting control load

Consider a model with a bi-sex, early-acting control load. As discussed above, this is obtained by setting 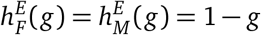 and 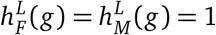 in Eqtn. (1) to get

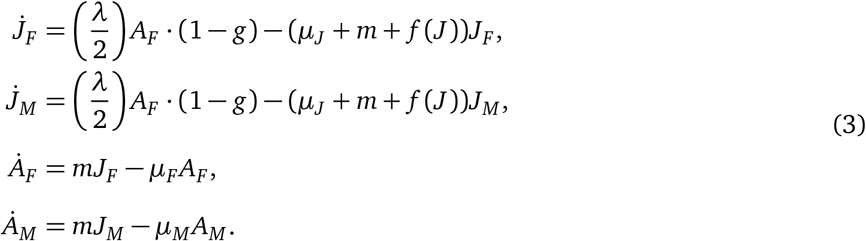

Solving for the equilibrium in the absence of control (*g* = 0) yields

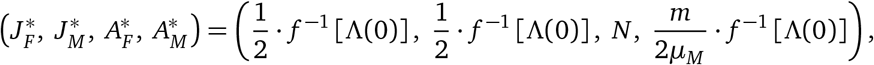

where *f* ^−1^ is the inverse function of *f*, and

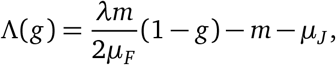

where we have used asterisks to denote populations at equilibrium. Here, *Λ*(0) is the per capita DD juvenile mortality rate in the absence of control. To allow for a fair comparison between systems, we adjust the density dependence parameter *α* to make the pre-control equilibrium equal for each choice of *f* (*J*).

The reproduction number, *R*_0_, determines the stability of the control equilibrium and is the average number of female individuals that live to adulthood produced by a single female throughout her lifetime when the population is small. This is the average total number of females produced by an adult female multiplied by the fraction of these eggs that survive to adulthood:

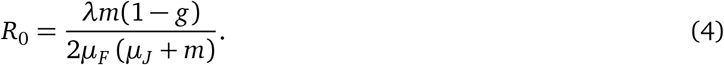

Note that DD juvenile mortality is negligible when the population is small, i.e. *f* (*J*) ≈ 0. In other words, the reproduction number is unaffected by density dependence. The 1 − *g* expression occurs in Eqtn. (4) because the female population is reduced by the control load. For a male-sex control load, eradication is impossible as we do not assume reproduction to be male-limited. In the absence of control (*g* = 0) and using the parameter values presented in Table 1, *R*_0_ = 33.14. We calculate the control load at which eradication is achievable for a population of *Ae. aegypti*. Stability analysis reveals that the extinction equilibrium is stable when *g* ≥ 1 − 1*/R*_0_ = 0.9698 and unstable otherwise. This means that a control load of approximately 97% or more is required to push the population to extinction.

We measure the success of population suppression by considering the adult female population at the control equilibrium. For the model in Eqtn. (3), the adult female population at the control equilibrium is

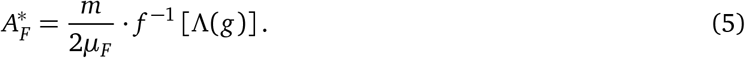

A female-sex, early-acting control load is modeled by setting 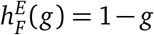 and 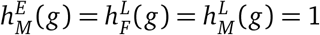 in Eqtn. (1). The adult female population at the control equilibrium in this case becomes

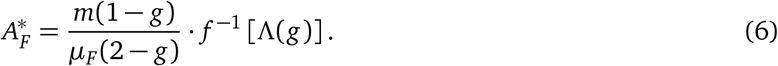

Comparing Eqtns. (5) and (6) and noting that (1 − *g*)*/*(2 − *g*) ≤ 1*/*2 for *g* ∈ [0, 1], we see that for an early-acting control load, female-sex targeting is more effective than bi-sex targeting. This is surprising because a bi-sex control targets both male and female juveniles and thus kills more organisms than a female-sex control. This happens because killing occurs prior to juvenile competition for a bi-sex control load, relaxing DD mortality more than for a female-sex control [68]. Fig. 2 shows the extent of population suppression for different strengths of juvenile DD mortality. The performance improvement of a female-sex over a bi-sex targeting control increases with the strength of density dependence. There is little performance difference in the system with weak density dependence, while the difference is substantial in the systems with stronger density dependence. Expressions for the remaining equilibrium populations are given in the Appendix.

**Figure 2:**
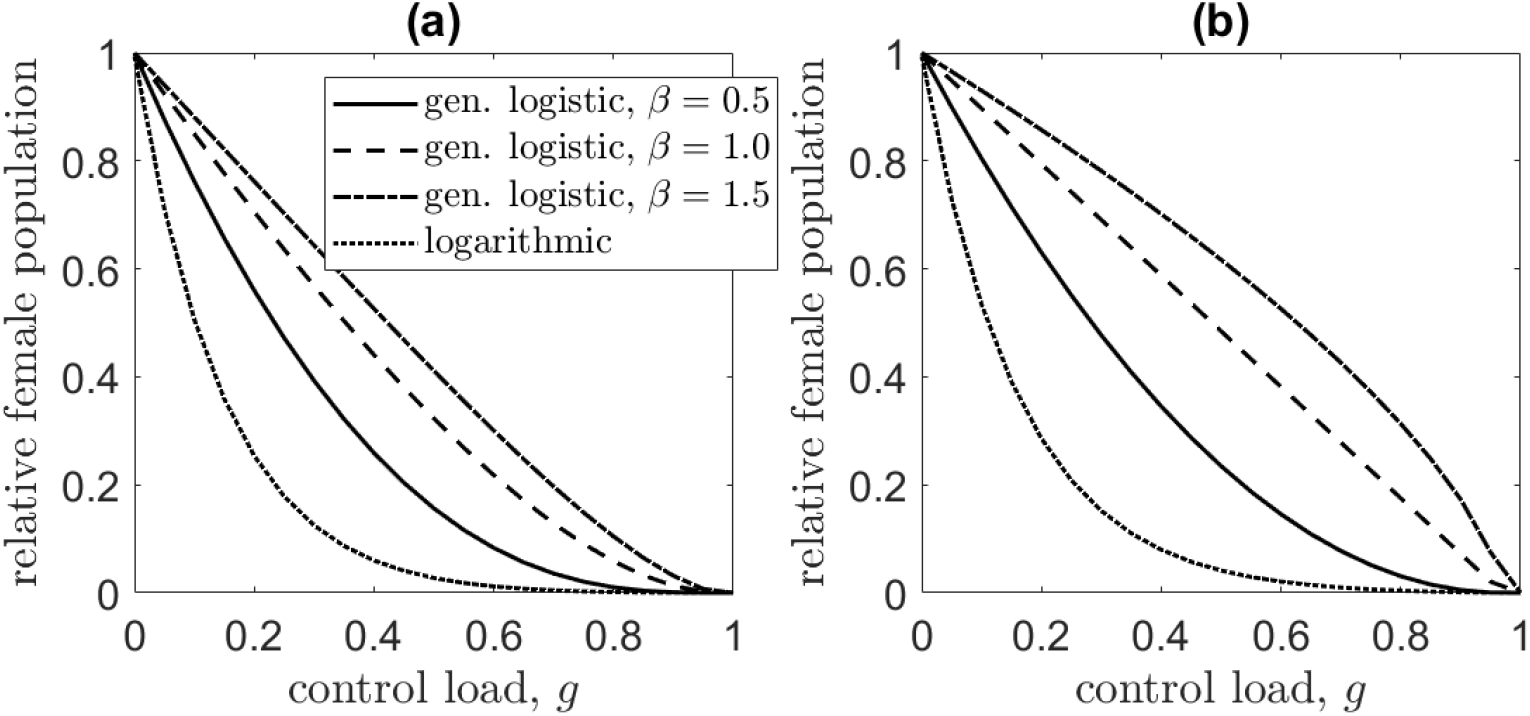
The adult female population at the control equilibrium for various levels of an early-acting control load when it is (a) female-sex and (b) bi-sex targeting. Line types indicate the per capita DD juvenile mortality function. The solid line denotes the generalized logistic case for *β* = 0.5, the dashed line corresponds to the *β* = 1 case, the dot-dashed line is *β* = 1.5, and the dotted line corresponds to the logarithmic case, *f* (*x*) = log 1 + (*αx*) ^*β*^.

The results presented in Fig. 2 indicate that population suppression is most successful in systems with weak density dependence. In our model, weak density dependence occurs for small *f* ^−1^(·), or those systems in which the per capita DD juvenile mortality is insensitive to changes in the population. In these systems, a perturbed population takes longer to reach equilibrium. Population suppression is less effective in systems with strong density dependence. The system with the weakest density dependence of those we consider, and consequently most vulnerable to population suppression, is that with the logarithmic form of juvenile DD mortality (see Fig. 1). In contrast, the population with the strongest density dependence—the generalized logistic system with *β* = 1.5—is most resistant to suppression. The performance difference between systems is sometimes dramatic, especially for low to intermediate bi-sex control loads. For a bi-sex control that kills 40% of juveniles (*g* = 0.4), for example, the generalized logistic system with *β* = 1.5 reduces the adult female population to only 70% relative to the pre-control equilibrium. For the logarithmic system, however, the population is reduced to 8% of its initial size. For an early-acting bi-sex control, performance differences are pronounced even for high control loads. This is not a feature of the early-acting female-sex targeting control.

### 4.2 Late-acting control load

A bi-sex, late-acting control load is obtained by setting 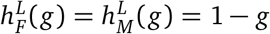 and 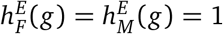 in Eqtn. (1):

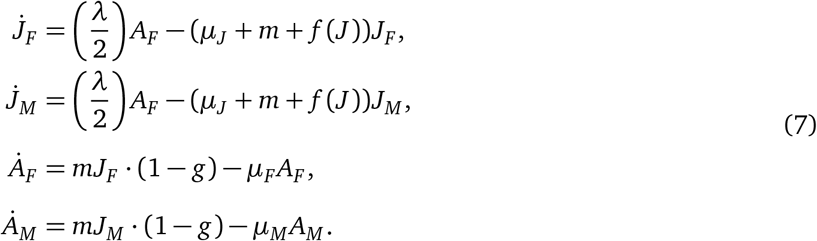

The reproduction number of the system with a late-acting control load (either female- or bi-sex) is the same as that of the system with an early-acting control load in Eqtn. (4). This is because the control load is imposed before females produce offspring. While the conditions for deterministic extinction are the same as in the previous subsection, the equilibrium values are not.

The adult female population at the control equilibrium is

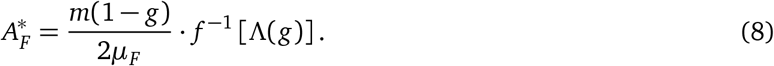

Since (1 − *g*)*/*2 ≤ (1 − *g*)*/*(2 − *g*) for *g* ∈ [0, 1], the late-acting control load is better at suppressing the population than either early-acting scenario considered in the previous subsection. This is because a late-acting control allows more individuals to contribute to DD juvenile mortality. The adult female population at the control equilibrium is plotted in Fig. 3 for different strengths of density dependence. The adult female population at the control equilibrium is equivalent between the systems with female- and bi-sex control loads. This happens because juvenile competition is unaffected by a late-acting control, and a reduced adult male population does not affect population dynamics.

**Figure 3:**
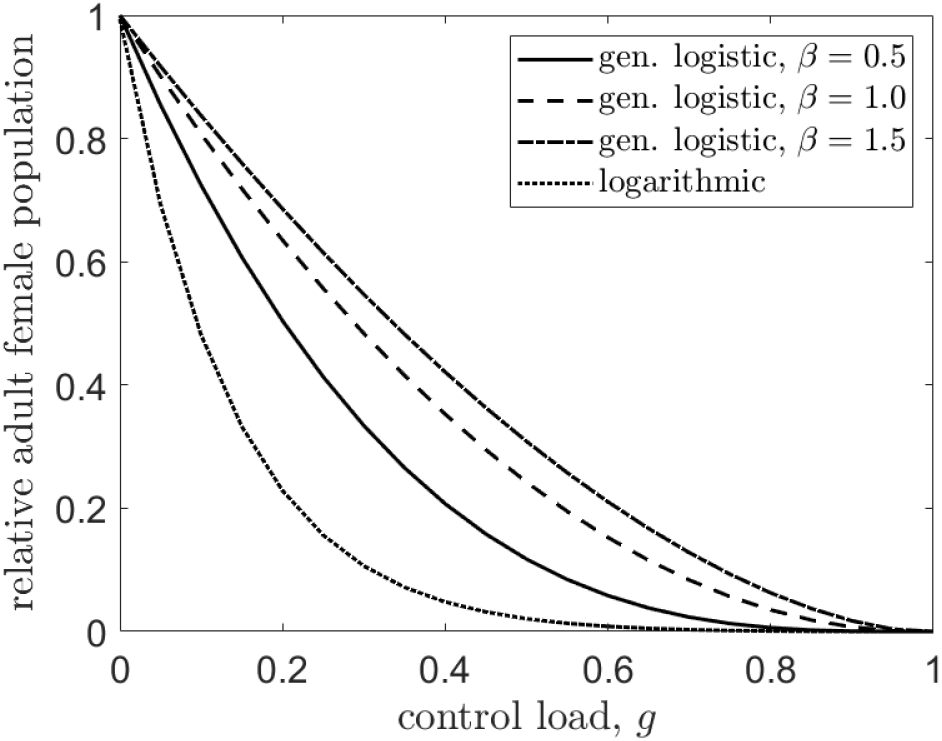
The adult female population at the control equilibrium relative to the pre-control population versus a late-acting control load. Results are equivalent for female- and bi-sex control loads. Line types indicate the functional form of per capita DD juvenile mortality. The solid line denotes the generalized logistic case for *β* = 0.5, the dashed line corresponds to the *β* = 1 case, the dot-dashed line is *β* = 1.5, and the dotted line corresponds to the logarithmic case, *f* (*x*) = log 1 + (*αx*) ^*β*^.

Referring to Fig. 3, performance varies most dramatically between the systems for intermediate control loads. For example, for a control that kills 50% of eclosing juveniles (*g* = 0.5), the generalized logistic system with *β* = 1.5 reduces the adult female population to approximately 31% relative to the pre-control equilibrium. For the logarithmic system, however, the population is nearly eradicated as it is reduced to 2% of its initial size. The success of population suppression is nigh indistinguishable for high control loads (*g* ≥ 0.8), in contrast to the results of the bi-sex early-acting control shown in Fig. 2(b). Thus, for a late-acting control load in a population with DD juvenile mortality, higher imposed mortalities overcome differences in density dependence strength in both bi- and female-sex targeting scenarios. This was not the case for the early-acting load, in which only the female-sex targeting control behaved in this way.

## 5 Results: Gene drive case studies

For each gene drive, performance is measured by the relative adult female population at the control equilibrium and, where appropriate, the invasion threshold required for successful introgression.

### 5.1 Homing gene drive

In our deterministic model, the construct becomes fixed in the population if the homing drive successfully invades. Therefore, to study the population size following successful control, one need only consider a population in which the transgene is fixed, or one in which all individuals have genotype *αα*. In this system, individuals incur a control load equal to their relative fitness, 1−*c*_*a*_. Using the notation of the general control load considered in the previous section, *g* = *c*_*a*_ by definition. Thus, a population in which the homing drive allele has become fixed is equivalent to the system with a late-acting, female-sex control load considered previously. (Recall that the results of female- and bi-sex control loads are equivalent for a general late-acting control.) The performance of a homing drive for a variable fitness cost, *c*_*a*_, is equal to that of the aforementioned system for variable control load shown in Figure 3.

### 5.2 One-locus engineered underdominance

We explore the performance of a one-locus EU drive for variable (ambient) fitness cost, *c*_*a*_, and for both bi- and female-sex targeting scenarios. The control equilibrium is determined numerically following transgenic release. Results are shown in Fig. 4. The one-locus EU gene drive is extremely effective at suppressing the population. Population suppression becomes more pronounced as the strength of density dependence weakens—see Figs. 4(a) and (b). In the system with the logarithmic form of DD juvenile mortality, for a modest fitness cost of *c*_*a*_ = 0.05, the gene drive crashes the female population to 1% of its original size. In contrast, for systems with stronger density dependence, population suppression is less impressive at only 9 − 26% relative to the pre-control population.

**Figure 4:**
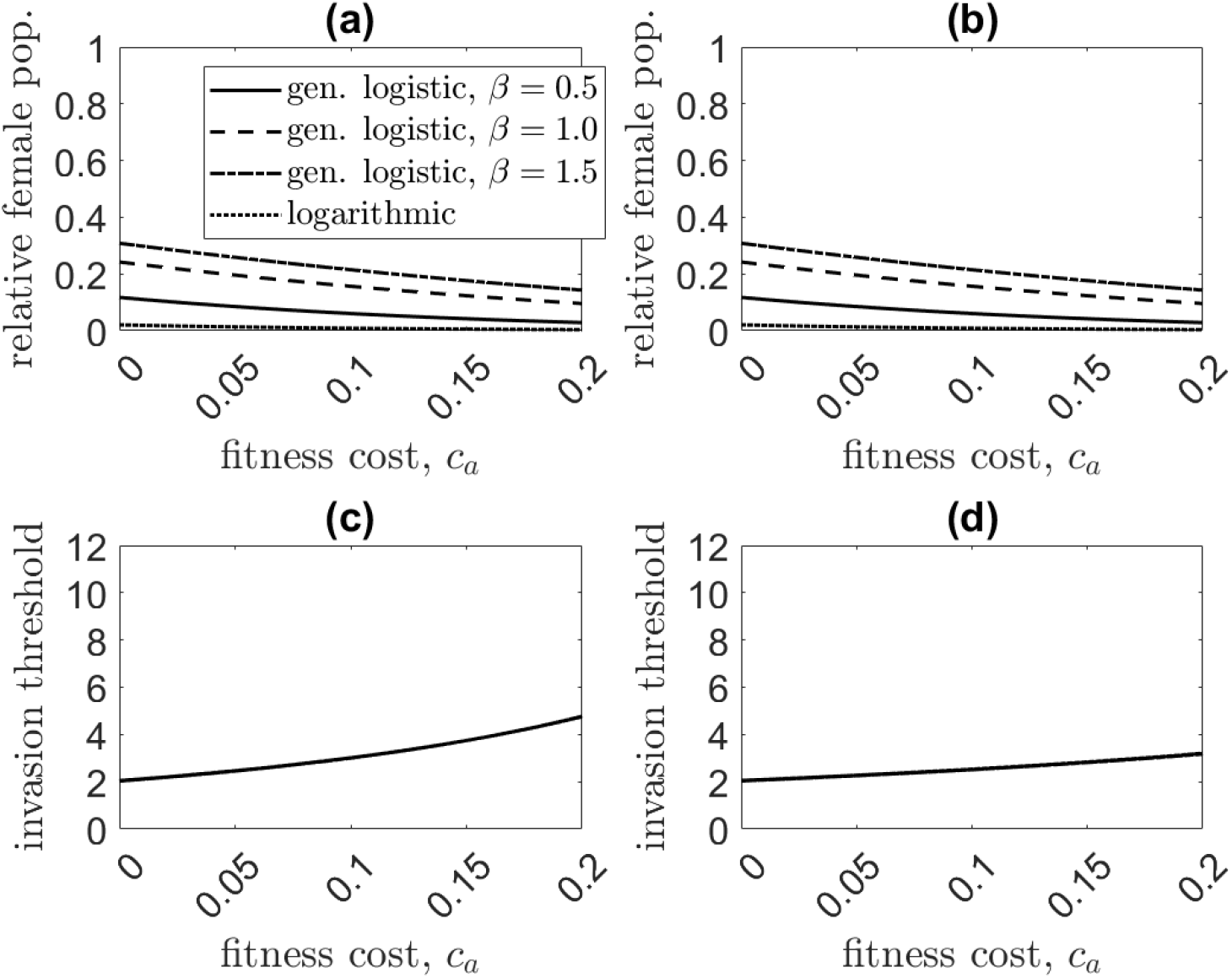
Comparison of the relative adult female population at the control equilibrium [(a) and (b)] and invasion threshold [(c) and (d)] for a female-sex (left column) and bi-sex targeting (right column) transgene for the one-locus EU drive as the ambient fitness cost (*c*_*a*_) is varied. The adult female population at equilibrium is equivalent between female- and bi-sex targeting cases, while the invasion threshold is lower for the latter. The density dependence functions studied are the generalized logistic case for *β* = 0.5 (solid line), *β* = 1 (dashed line), *β* = 1.5 (dot-dashed line), and the logarithmic case *f* (*X*) = log 1 + (*αX*) ^*β*^ (dotted line).

Gene drive performance differs slightly between female- and bi-sex control loads. The adult female population at the control equilibrium is equivalent between systems, but the bi-sex targeting transgene invades at slightly lower release ratios than the female-sex targeting transgene, as indicated by a lower invasion thresh-old. This result is not obvious, as the control load in the bi-sex system is imposed on both sexes and thus would reasonably be expected to require larger releases for invasion. However, for the female-sex targeting system, males can carry wild-type alleles with little consequence, effectively serving as reservoirs for wild-type alleles [24, 68]. A higher release ratio is then required for invasion and, additionally, a lower transgene frequency at equilibrium is achieved. For either scenario, the invasion thresholds do not differ substantially with the strength of density dependence.

### 5.3 Two-locus engineered underdominance

The analysis of the previous subsection is repeated for the two-locus case, with an important difference being that a two-locus EU drive can use male-only releases to introduce the transgene. The results of our analysis are shown in Fig. 5. The two-locus EU gene drive is less successful than the one-locus case in both population suppression and invasion threshold. Higher invasion thresholds were observed for the two-locus case, especially for the female-sex targeting transgene. However, this is mostly due to our using a male-only release for the two-locus EU drive, versus a bi-sex release for the one-locus EU drive. If a bi-sex release is used for the two-locus EU drive, the invasion threshold is lower than that of the one-locus case (results not shown). As in the one-locus case, the bi-sex targeting system required smaller release ratios to invade than the female-sex targeting system. This happens for the same reasons described above: males are still viable if they carry a toxin allele without a complementary antidote, and can thus harbor wild-type alleles longer than if the construct was bi-sex targeting [24]. The female-sex targeting system performed slightly better at reducing the adult female population at equilibrium than the bi-sex system (see Figs. 5(a) and (b)). This difference is only apparent for fitness costs *c*_*a*_ *>* 0.04.

**Figure 5:**
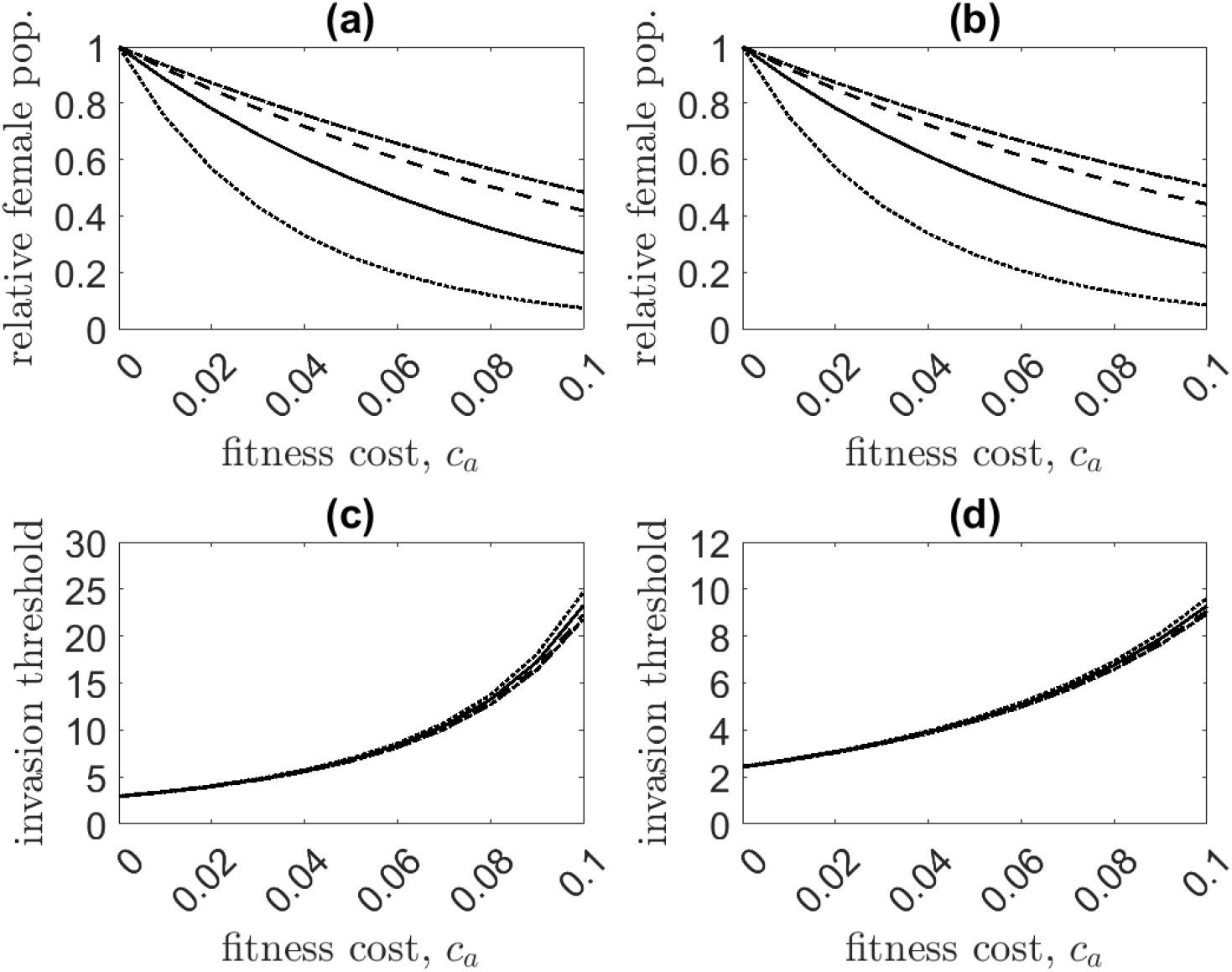
Comparison of the relative adult female population at the control equilibrium [(a) and (b)] and invasion threshold [(c) and (d)] for a female-sex (left column) and bi-sex targeting (right column) transgene for the two-locus EU gene drive. We assume that the toxin is lethal without the complementary allele (*c*_*t*_ = 1). The density dependence functions studied are the generalized logistic case for *β* = 0.5 (solid line), *β* = 1 (dashed line), *β* = 1.5 (dot-dashed line), and the logarithmic case *f* (*X*) = log 1 + (*αX*) ^*β*^ (dotted line).

Population suppression is most effective in the system with the weakest density dependence and progressively worsens with increasing strength of density dependence. For a small fitness cost of *c*_*a*_ = 0.07, the invasion threshold of the bi-sex lethal system is approximately 5.8, while for the female-sex lethal system, it is slightly over 10. For comparison, a recent field application of SIT against *Ae. aegypti* used a release ratio of 6.4 [23]. Population suppression is comparable between the bi- and female-sex targeting payloads but differs greatly based on the strength of density dependence, as expected from the results of the previous section. For the same ambient fitness cost of *c*_*a*_ = 0.07, for the logarithmic form of density dependence, the population is suppressed to approximately 15% of its size relative to pre-control counts. In the systems with logistic density dependence, the performance is less impressive at around 40 − 60% relative to the pre-control population.

Interestingly, the invasion threshold varies with the strength of density dependence in Figs. 5(c) and (d). This suggests that the strength of density dependence also influences the number of released transgenic insects required for invasion—populations with weak density dependence require slightly higher releases than those with strong density dependence. Closer investigation reveals that these differences are caused by a combination of factors, including the strength of density dependence and assumptions of adult lifespan for each sex. The rate at which progeny emerge to mate with the released males and produce drive-homozygous progeny is determined by the strength of density dependence, as this controls how quickly a population responds to perturbations. In the time it takes for this to happen, however, the released males die at per capita rate *µ*_*M*_. When the first heterozygote progeny eclose, they encounter a smaller transgenic male population than at the time of release. Adult lifespans also determine the ratio of males to females at equilibrium. If females live longer than males (*µ*_*F*_ *< µ*_*M*_), as is the case with *Ae. aegypti* [21, 50, 61], wild-type alleles persist longer in the female population. If the lifespans between sexes are equal (*µ*_*M*_ = *µ*_*F*_), alleles persist for equally long in either sex, and the eclosion rate (controlled by the strength of density dependence) matters less. While we have discussed male-only releases, this is true for any single-sex release. To reduce the impact of density dependence, multiple and/or bi-sex releases can be employed. Bi-sex releases ensure that drive homozygous progeny are produced in the first generation following release, hence why invasion threshold differences are less apparent for the one-locus EU gene drive.

## 6 Discussion

Here we have used mathematical models with sex, age, and genetic structure to study the relationship between the strength of density dependence and population suppression. We first considered performance differences as the timing of a general control was varied. This was either early- or late-acting, distinguished by whether a control load was imposed before or after DD juvenile mortality, respectively. Timing the control so that it exploits DD factors can be beneficial to suppression. Certain pests, such as *Ae. aegypti* [2, 64, 70, 72] and other mosquito species [42, 71, 80], experience DD juvenile mortality. If suppression occurs after DD competition, individuals destined to be killed still compete and contribute to reduced juvenile survival [24, 56, 77]. As the example of *Ae. aegypti* demonstrates, leveraging the ecology of a pest species is critical in developing an effective control program [53]; when done incorrectly, it can have the paradoxical consequence of increasing the target population [1, 47]. However, note that the continuous-time models used in this study cannot exhibit overcompensatory density regulation.

For a general control load, including the gene drives we studied, we find that a female-sex targeting system is equally or more successful at reducing the female population at equilibrium than a bi-sex targeting system. The difference in performance, if any, is more substantial for stronger density dependence; that is, in populations with weak density dependence, there is little difference between female- and bi-sex targeting systems for an early-acting control load. The advantage of a female-sex targeting control over one that targets both sexes increases with the strength of density dependence. For a general late-acting control load in our model, there is no difference between female- and bi-sex targeting systems in reducing the female population at equilibrium. Female-sex control loads are typically preferred as it tends to be the females of a pest species that are the most injurious [55]. In the case of medically important mosquitoes, only females blood feed and thus transmit pathogens.

The baseline model used to study a general control load was then extended to simulate the performance of three specific genetic biocontrol strategies: a simple homing gene drive, and a one- and two-locus EU drive. All these gene drives were assumed to impose a late-acting control load. The performance of the homing gene drive was identical to that of a general late-acting control load—there was no difference in the female control equilibrium between female- and bi-sex targeting transgenes. For a one-locus EU drive, population suppression of the female population at the control equilibrium was incredibly successful and equivalent between female- and bi-sex control loads. However, successful invasion required a bi-sex release since we assumed the toxin allele renders a transgenic organism nonviable if it lacks the complementary antidote allele. A bi-sex targeting transgene was able to invade the population at slightly lower release ratios than a transgene targeting only females. This is likely a result of male heterozygotes being able to carry transgenes without ill effect in the case of a female-sex targeting transgene [24, 68]. The difference in the invasion threshold was only moderate. The same phenomenon occurs in the two-locus EU case, explained by similar reasoning.

Unlike the one-locus case, for the two-locus EU drive, a female-sex targeting transgene performed slightly better at reducing the female population than a bi-sex targeting transgene. Gentile et al. (2015) observed the opposite trend, albeit in the case of transgenic SIT: late-acting bi-sex killing was slightly more effective than late-acting female-sex killing at reducing the adult female population. Vella et al. (2021) clarify that the performance of either control relative to each other depends on the manner of transgenic releases [68]. Despite the improved suppression performance, we found that a female-sex targeting two-locus EU drive required larger releases for invasion to occur than a bi-sex system, agreeing with the findings of [68] and [34]. Interestingly, we also found that the strength of density dependence, as well as assumptions of adult lifespan, can influence the invasion threshold of a gene drive. This occurred for single-sex releases of a two-locus EU drive. A larger release ratio for successful introgression may be needed when density dependence is weak and there is a large difference in lifespan between sexes. This deficiency can be mitigated by multiple or even bi-sex releases, depending on the pest species.

Our study has several limitations. We have not considered the contribution of space to our results. Control program success is affected by spatial heterogeneity, non-uniform pest distributions, and proximal mating [19]. All of these factors can hinder the success of population suppression in practice. Furthermore, the spatial distribution of a pest population might itself depend on density, such as in response to parasitoid abundance [16]. In the interest of keeping our model as simple as possible, we consider only DD juvenile mortality. For *Ae. aegypti*, both development time and adult body size are density-dependent [2, 49, 70, 71, 72, 80]. While these factors were not included in our analysis, it is reasonable to expect these would exacerbate differences in control performance between systems. We do not account for Allee effects in our model. For sufficiently small populations, successful mate finding can frustrate population growth, as has been studied in other population dynamics models of *Ae. aegypti* [51] and in modeling studies of the genetic control of mice [8]. Small populations—such as might be caused by seasonality or population control—would also incur a fitness cost due to inbreeding depression. The work of [60] found that *Ae. aegypti* is severely impacted by inbreeding depression: survival rates, fecundity, egg hatch proportions, and male mating success are reduced when compared to larger populations. Egg-laying site choice behavior is also impacted by conspecific density [17]. Reducing *Ae. aegypti* numbers can swing the competitive edge to favor that of cohabitant mosquito species, such as *Ae. albopictus* [12]. These considerations paint an encouraging picture of the level of suppression required to achieve eradication compared to that predicted by our deterministic analysis. Indeed, population eradication need not require that every individual pest be killed [40]. Following similar logic, eradicating a pest population is unnecessary to reduce its burden to tolerable levels, including the burden of pest-vectored disease. The field study in [79] found an appreciable reduction in malaria burden was achieved with a 70% reduction in the mosquito population. The authors of [41] found that dengue cases within the field study area fell by over 70% following an 80% reduction in the number of containers testing positive for the presence of *Ae. aegypti*. However, it is important when implementing population control to prevent pest populations from returning to pre-control levels, as this could have severe consequences. This is especially true for vectors of harmful diseases as transient population control leads to the accumulation of susceptibles [31].

We opted to employ a general density dependence function as there are few quantitative results on density-dependent dynamics in wild populations of *Ae. aegypti* [20, 38, 64]. Further study in this direction is critical to accurately inform models of mosquito population control. This is not unique to mosquitoes. As we have shown, population suppression efficacy can vary markedly depending on assumptions surrounding the strength of density dependence, which can have important consequences on the outcome. While our model lacks a settled mathematical formulation of juvenile density dependence, we have compensated for this— and consequently made our work more generalizable—by studying a variety of scenarios. Populations with strong density dependence are more resilient to population control and can return to pre-control equilibrium quickly following a reduction. From studies of mosquito populations pre- and post-control, this appears to be the case for *Ae. aegypti* [36, 57, 73]. The authors of [73] evaluated the effect of larvicide and adulticide applications on *Ae. aegypti* populations. After the control had ceased and the mosquito population had been reduced by nearly 88%, it took only 9 weeks (approximately 8-9 generations) for it to return to pre-treatment levels. For pest species that exhibit weak density dependence (i.e., those species where population regulation is slow), population suppression is more effective.

Our findings demonstrate that the strength of density dependence in a pest population can substantially alter control program performance. Furthermore, we have shown that taking into account the natural regulatory factors of a pest population can increase the performance of control measures, sometimes dramatically. Ignoring these factors risks control failure or even worsening pre-control conditions. Beyond practical shortcomings, misquantifying density dependence in a mathematical model can lead to inaccurate control predictions. The density dependence of a pest must be understood before implementing effective population control.

## Acknowledgments

The authors are grateful for Drs. Fred Gould and Jen Baltzegar, as well as Casey O’Brien, for their valuable feedback and advice while preparing this manuscript. CDB was supported by the NSF under award DGE-2137100. ALL and CDB received support from NIH grant R01-AI139085.

## 7 Appendix

### 7.1 Clarification of model formulation in Bellows [7]

The logarithmic form of density dependence we consider in the main text, *f* (*x*) = ln 1 + (*αx*) ^*β*^, is discussed by Bellows [7] and originally proposed in [63]. Many classic studies of the dynamical impact of density dependence use discrete time, non-overlapping generation, models [29, 43, 44]. Bellows [7] considered the correspondence between continuous and discrete time descriptions of density dependence and constructed a table summarizing both forms (see Table 1 in [7]). Bellows employed the general differential equation

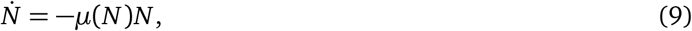

where *N* is the population density and *µ*(*N*) is the per-capita mortality rate. Population survival over some time was derived by integrating Eqtn. (9) over a generation (taken to be one time unit). In general, this integration cannot be performed analytically. In order to allow derivation of a closed-form discrete-time model, Bellows employed a constant per-capita mortality assumption, approximating the function *µ*(*N*) by its *initial value, µ*(*N*_0_). Under this approximation, he showed that the logarithmic form for mortality gives rise to the following discrete-time model

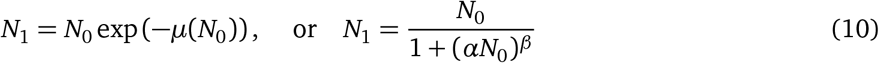

when *µ*(*N*_0_) = ln 1 + (*αN*_0_)^*β*^. This model was familiar in the literature and exhibits overcompensation. This result has led to the claim that the logarithmic form of density dependence gives rise to overcompensation in continuous time models.

Bellows’ analysis (and hence the correspondence between the two models) relies on his assumption of a constant per-capita mortality rate. However, even over small time intervals, assuming a fixed density of organisms is not a good approximation of the continuous-time system. Numerical integration of Eqtn. (9) shows that there is a qualitative difference between this solution and the solution in Eqtn. (10) arrived at under Bellows’ simplifying assumption (see Figure 6).

**Figure 6:**
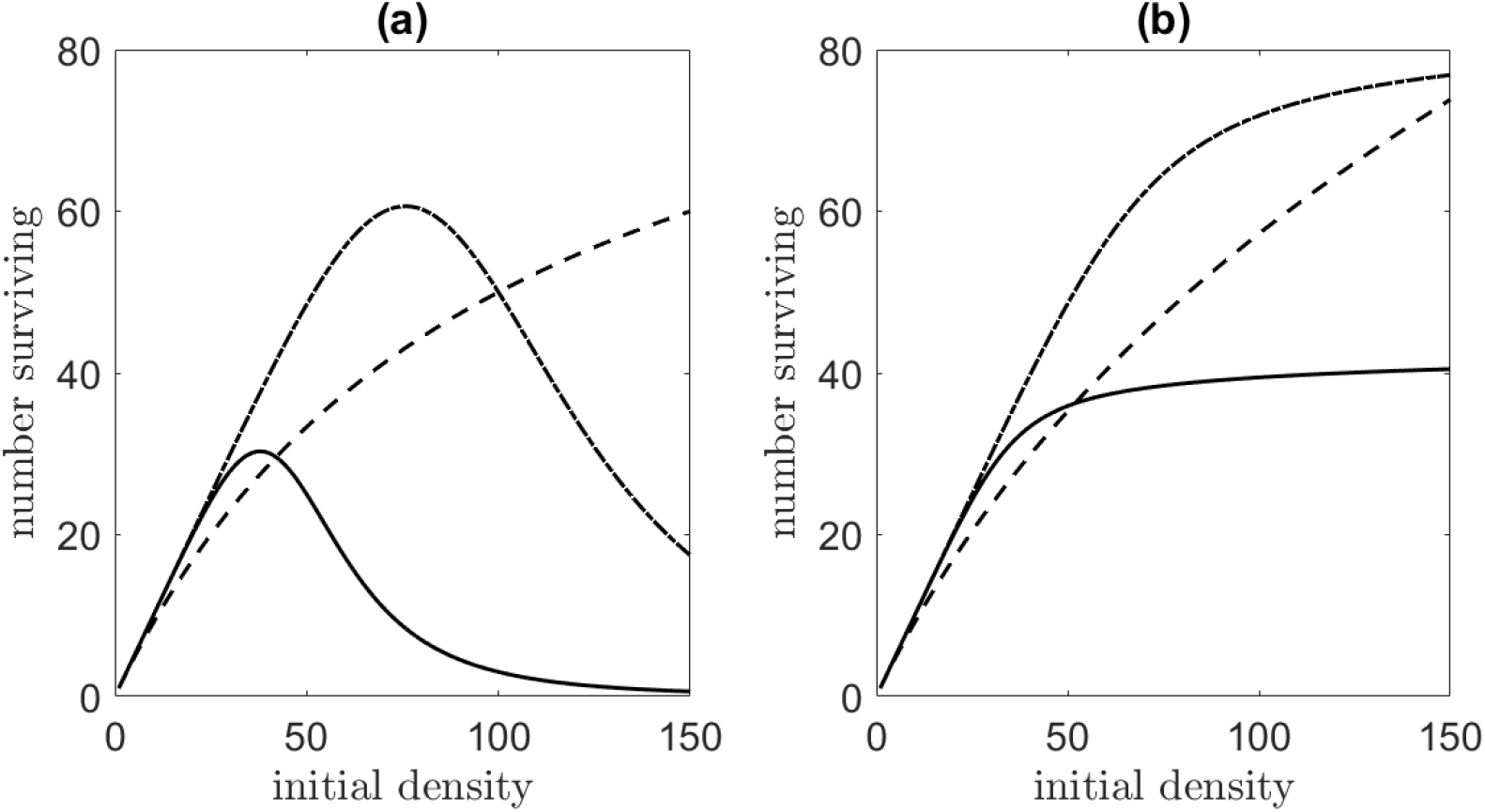
A comparison of the number of surviving organisms at *t* = 1 for the (a) discrete system in Eqtn. (10) and the (b) continuous-time system in Eqtn. (9) for *µ*(*N*) = log 1 + (*αN*) ^*β*^ for variable initial densities. The different parameter values used are: *α* = 0.01, *β* = 1 (dashed), *α* = 0.01, *β* = 5 (dot-dashed), and *α* = 0.02, *β* = 5 (solid). The discrete system in (a) exhibits overcompensatory density dependence, or scramble competition, while the continuous-time differential equation in (b) does not.

For the discrete formulation in Eqtn. (10), it was shown that individual survival decreases for larger initial densities (i.e., overcompensation or scramble competition) when *β* ≫ 1. This is not a property of the continuous-time formulation in Eqtn. (9). We show *N*_1_ against *N*_0_ for Eqtn. (9) with *µ*(*N*) = log 1 + (*αN*) ^*β*^ alongside the discrete system in Equation (10) for different values of *α* and *β* in Fig. 6. Note that the curves for the discrete system are exactly those presented in Fig. 6(a) in [7]. For the continuous-time system, it is never the case that the number of survivors is maximized at an intermediate initial density. Indeed, a unimodal discrete-time map can never result from a mortality process modeled by a single ordinary differential equation [26]. However, such behavior can occur in a system of differential equations if a time delay is included (e.g., [4, 25]).

### 7.2 Equilibria for the system with an early-acting control load

The equilibrium of the wild-type populations with an imposed control load is calculated for each lethality scenario (male-, female-, and bi-sex targeting) when the per capita density-dependent juvenile mortality is given by the general function *f* (*J*). We require that *f* (*x*) is invertible over the domain [0, ∞). We let

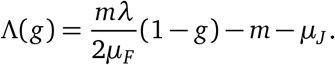

In the equilibria provided below, *f* ^−1^ denotes the inverse function of *f*.

#### Bi-sex lethality

When there is bi-sex lethality, the control load is applied to both sexes. The equilibrium is given by

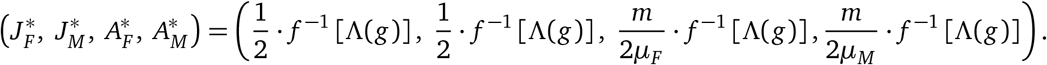

#### Female-sex lethality

In the case of female-sex lethality, the control load is imposed only on the female population. The equilibrium is

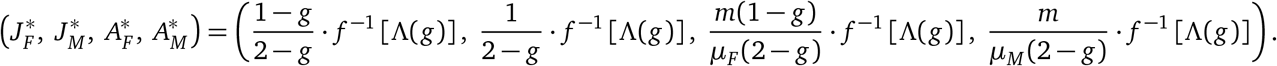

#### Male-sex lethality

In the case of male-sex lethality, the control load is imposed only on the male population. The equilibrium is

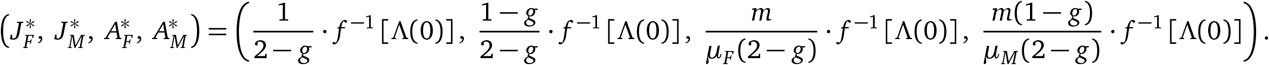

### 7.3 Equilibria for the system with a late-acting control load

Equilibria of the wild-type populations with a late-acting control load are calculated for each lethality scenario (male-, female-, and bi-sex targeting). We let

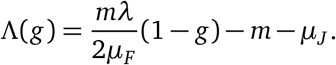

In the equilibria provided below, *f* ^−1^ denotes the inverse function of *f*.

#### Bi-sex lethality

When there is bi-sex lethality, the control load is applied to both sexes. The equilibrium for a late-acting, bi-sex lethal control load is

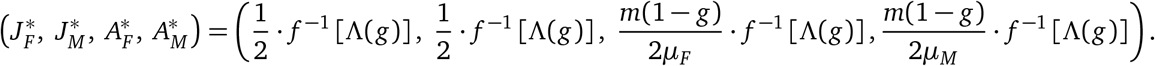

#### Female-sex lethality

In the case of female-sex lethality, the control load is imposed only on the female population. The equilibrium is

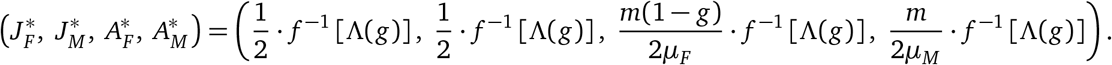

#### Male-sex lethality

In the case of male-sex lethality, the control load is imposed only on the male population. The equilibrium is

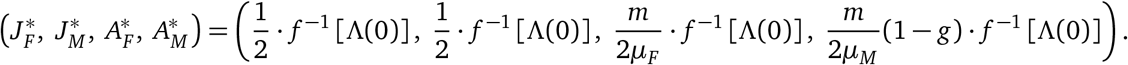

